# Bridging Computational Neuroscience and Machine Learning on Non-Stationary Multi-Armed Bandits

**DOI:** 10.1101/117598

**Authors:** George Velentzas, Costas Tzafestas, Mehdi Khamassi

## Abstract

Fast adaptation to changes in the environment requires both natural and artificial agents to be able to dynamically tune an exploration-exploitation trade-off during learning. This trade-off usually determines a fixed proportion of exploitative choices (i.e. choice of the action that subjectively appears as best at a given moment) relative to exploratory choices (i.e. testing other actions that now appear worst but may turn out promising later). The problem of finding an efficient exploration-exploitation trade-off has been well studied both in the Machine Learning and Computational Neuroscience fields. Rather than using a fixed proportion, non-stationary multi-armed bandit methods in the former have proven that principles such as exploring actions that have not been tested for a long time can lead to performance closer to optimal - bounded regret. In parallel, researches in the latter have investigated solutions such as progressively increasing exploitation in response to improvements of performance, transiently increasing exploration in response to drops in average performance, or attributing exploration bonuses specifically to actions associated with high uncertainty in order to gain information when performing these actions. In this work, we first try to bridge some of these different methods from the two research fields by rewriting their decision process with a common formalism. We then show numerical simulations of a hybrid algorithm combining bio-inspired meta-learning, kalman filter and exploration bonuses compared to several state-of-the-art alternatives on a set of non-stationary stochastic multi-armed bandit tasks. While we find that different methods are appropriate in different scenarios, the hybrid algorithm displays a good combination of advantages from different methods and outperforms these methods in the studied scenarios.

## 1 Introduction

Selecting an action from a number of distinctive alternatives in unknown environments, where each action may return a reward in a stochastic manner, can be a challenging problem when the environment is non-stationary (i.e the rewards are changing in time). The adaptive learning strategy of the decision maker, especially when facing abrupt changes, requires a sophisticated method for the well known exploration-exploitation trade-off. Modeling the problem for implementing intelligent decision-makers is a subject mainly studied from the machine learning field, while the computational neuroscience field is concerned about finding models that fit human’s or animal’s behavior efficiently. In the former the task is to find a decision-making algorithm whose actions achieve optimal cumulative regret - the regret being defined as the difference between optimal performance and the accumulated reward. Solutions to the stationary problem have been studied thoroughly for more than a decade, but the non-stationary approach was a more recent subject of research [1,2], while its recent recapturing of attention is particularly due to its characteristic as a keystone in reinforcement learning. In parallel, computational neuroscience researches have also progressed in combining methods from reinforcement learning with exploration bonuses [3] and sometimes change-point detectors [4] to tackle the problem of fitting a human’s or an animal’s action decisions on bandit tasks with altering rewards in time.

With the dual goal of conceiving adaptive machines with cognitive components (e.g for human-robot interaction environments) and of generating new experimental predictions for neuroscience, we explored the impact of integrating approaches from both fields at the lower level of analysis for a simple artificial decision-maker. We tested the performance of several alternative algorithms on a set of different non-stationary bandit setups where we variated the most crucial components of a stochastic and changing environment. Here we present empirical evidence that our hybrid approach outperforms some state-of-the-art alternatives, rising questions about how the integration of neuroscience-based models and machine learning-models could have a positive impact on performance.

## 2 Problem Formulation and Algorithms

Stochastic multi armed-bandits can be considered as having a set of arms *𝒦* = {1, 2,…, *K*} of a casino machine, where a decision-maker chooses an action *a* ∈ *𝒦* at every timestep *t* ∈ *𝒯* = {1,2,…, *T*} of its lifespan *T* (also called the time horizon). He then receives a reward *r*_*t*_(*a*) with probability *p_t_*(*a*) and zero otherwise. While interested in arm’s expected value of reward, without loss of generality, we can assume Bernoulli arms where ∀(*a*, *t*) ∈ *𝒦* × *𝒯* the rewards are binary, *r_t_*(*a*) ∈ {0,1}. By choosing Bernoulli arms, the expected value 𝔼[*r_t_*(*a*)] for every arm reward is equal to the probability *p_t_*(*a*). When the environment is stationary, *p_t_*_+1_(*a*) = *p_t_*(*a*) for all time steps. For non-stationary environments the above rule does not stand. Specifically in drifting environments |*p_t_*_+1_(*a*) − *p_t_*(*a*)| < *ϵ*, where *ϵ* is a small constant, while in abruptly changing environments ∃*t*: |*p_t_*_+1_(*a*) − *p_t_*(*a*)| > *δ*, where *δ* is a sufficiently large value in terms of probabilities. Both *ϵ* and *δ* can be used as a measure of the environment dynamics. Denoting *a*\* the optimal arm to choose, the regret at every time step is then defined as 𝔼[*r_t_*(*a*^*^) − *r_t_*(*a*)].

A first important note is that in most bandit methods, from either machine learning or computational neuroscience, the action *a_t_* to be taken at the time episode *t* can be chosen after comparing between all actions a combination of an empirical measure of each action’s value *μ_t_*(*a*) and some measure of uncertainty for this value *σ_t_*(*a*), according to the following general equation:

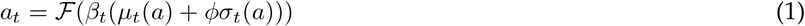

In the classical family of bandit methods initially developed in machine learning - called *UCB* for Upper Confidence Bound [5] -, function *ℱ* is an argmax, parameters *β* and *ϕ* are fixed to 1, and *μ_t_*(*a*), *σ_t_*(*a*) are empirically estimating the average reward yielded by action *a* and the uncertainty associated to this estimate. Then the method differ in the way these estimates are updated, using for instance a *Sliding Window* (SW-UCB) or *Discounted* estimates (D-UCB) [1]. In a more recent work, the *Kalman Filter Multi-Armed Normal Bandit* (KF-MANB) proposed in [2] is also another empirically optimal algorithm using an observational noise parameter *σ_ob_* and a transitional noise parameter *σ_tr_* to estimate *μ_t_*(*a*) and *σ_t_*(*a*) at each timestep, which are then used as the parameters of a normal distribution *𝒩*(*μ*_*t*_(*a*), *σ_t_*(*a*)) for each arm. Then a sample *s_a_* from each distribution is taken, and the action choices are made according the rule *a_t_* = arg max_*a*_ (*s_a_*).

In computational neuroscience models for bandit tasks [3,6], function *ℱ* is a Boltzmann softmax followed by sampling from the resulting action probability distribution, parameters *β* and *ϕ* are either fixed or estimated from each human subject’s behavior during the task, *μ_t_*(*a*) is the *Q-value* of action *a* learned through reinforcement learning - most often with a discounted factor *γ* = 0 because the task is not sequential - and *σ_t_*(*a*) is an exploration bonus. In some models, *μ_t_*(*a*) can be transiently adapted faster through the use of change point detectors which identify abrupt task shifts that produce a higher noise than the expected observational noise [4]. Interestingly, this has also been recently applied to bandit methods in Machine Learning (*e.g.* the *Adapt-Eve* algorithm [7]). Finally, some models propose to dynamically tune through *meta-learning* the algorithm’s parameters as a function of variations of performance [8,9]: for instance, increasing the inverse temperature *β_t_* in the softmax function to promote exploitation when the average performance improves, and decreasing it to promote exploration when performance drops indicate abrupt changes in the task.

In previous work, we have applied this meta-learning principle in an algorithm here referred to as *MLB* to dynamically tune *β_t_* in a simple multi-armed bandit scenario involving the interaction between a simulated human and a robot [10]. Formally, function *ℱ* is a Boltzmann softmax with parameter *ϕ* = 0 (*i.e.* no uncertainty term is used as exploration bonus here). The immediate reward *r_t_* received from action *a = a_t_* is then used to update the estimated action’s expected value *μ_t_*(*a_t_*) of the action taken, using the update rule of a low-pass filter, as *Q*_*t*+1_(*a*) = (1 − *α_Q_*)*Q_t_*(*a*) + *α_Q_r_t_*. Furthermore, following [8], a similar update rule applies to estimate mid-term and long-term reward running averages according to:

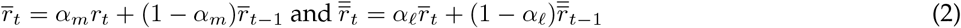

where *α_m_* and *α_ℓ_* are two different learning rates. *β_t_* is then updated as

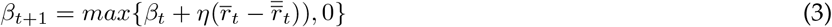

With the above update rules, *β_t_* increases when the mid-term reward is greater than the long-term. One can view this as an evidence that the recent actions are more optimal, therefore exploitation may be increased. In the opposite case when *r̅_t_* < *r̿_t_*, the recent actions denote that the present policy has lower performance in getting rewards compared to the past, therefore exploration should be encouraged by reducing *β_t_*.

## 3 Hybrid Meta-learning with Kalman Filters - MLB-KF

After showing a common way to formalize bandit methods in machine learning and computational neuroscience, the objective of the present paper is to illustrate how a hybrid method combining ideas from the two fields that we have recently proposed in [11] can yield a promising performance in a variety of non-stationary multi-armed bandit tasks. The hybrid model integrates the sibling kalman filters of KF-MANB and the core of MLB. To do so, action values learned through reinforcement learning in *MLB* are here simply replaced within the softmax function with a linear combination (with *ϕ* > 0) of the mean *μ_t_*(*a*) and the standard deviation *σ_t_*(*a*) of each arm’s distribution according to the sibling kalman filters of KF-MANB. The algorithm also uses two constants 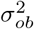 and 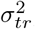, which relate to the stochastic part and the nonstationary part of the environment respectively. The values for means *μ*_1_(*a*), variances 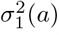, and *β_t_* are initialized. Following [2], but writing equations down on a different way, the reward *r_t_*(*a*), for *a = a_t_*, is then used to update the parameters of the respective arm with

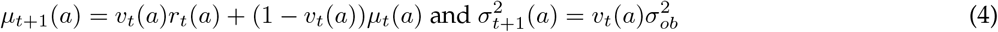

while for arms *a ≠ a_t_* the means *μ*_*t*+1_(*a*) maintain their previous value, and a transitional variance is added to 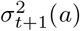 in order to enhance exploration of non-chosen arms as shown below

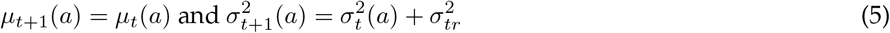

In the above equations *v_t_*(*a*) are dynamically tuned learning rates for each arm, which may mirror some adaptation procedures of learning rate studied in a computational neuroscience bandit task (*e.g.* [12]). Here, for KF-MANB, it follows:

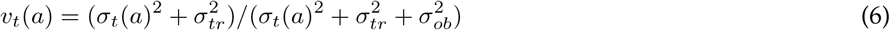

The mid-and long-term rewards are also calculated like in [8] and used to update the inverse temperature parameter *β_t_* according to the update rule of MLB [10]. When the uncertainty of an arm’s action value increases, the respective arm should be explored more. However when the environment suggests that no evidence of change is observed, KF-MANB suffers from exploration since a transitional variance is always added to the non-pulled arms. This can also be seen in Fig.1 (right) of simulations section. MLB is also expected to display large regret at cases where a non-optimal arm becomes optimal after long periods of stationarity, since the "rising" arm may never be chosen. The above implementation is a trade-off between these two cases, as we empirically present evidence that it incorporates both the benefits of MLB’s fast adaptation and small regret on many test cases, as also the robustness and good performance of KF-MANB.

**Figure 1:**
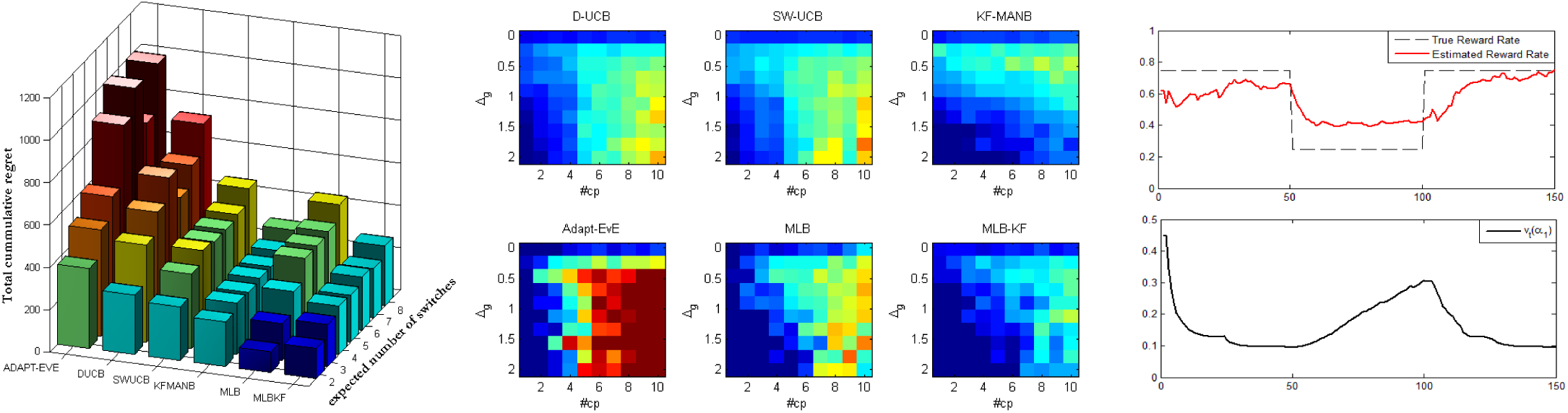
Left: total regret for different values of expected number of change points *h*, middle: total cumulative regret for different gaps ∆_*g*_ and number of switches *#cp* (thresholded for low and high values), right: estimated action value *μ*(*a*_1_) vs true value (top) and parameter *v_t_*(*a*_1_) of KF-MANB for a two-armed bandit case (bottom)

~~~
**Algorithm 1** Hybrid MLB-KF
1: Choose parameters *α_m_, α_ℓ_, η, ϕ, σ_ob_, σ_tr_*
2: Initialize *β_1_*, and *μ_1_*(*a*), σ_1_(*a*) ∀*a ∈ 𝒦*
3: **for** *t* = 1, 2,…, *T* **do**
4:   select action *a_t_* ∈ *𝒦* from distribution of (1) with <u>s</u>oftmax
5:   observe the immediate reward *r_t_* and update *r̅_t_*, *r̿_t_* with (2)
6:   for *a* = *a_t_* update *μ_t+1_*(*a*), 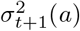 with (4),(6)
7:   for all arms *a* ≠ *a_t_* update *μ_t+1_*(*a*), 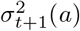 with (5)
8:   update *β_t+1_* with (3)
9: **end for**
~~~

## 4 Numerical Simulations

Testing an algorithm on a scenario that was also used in order to tune its parameters, should restrict its evaluation of performance to concern only this specific environment. Here we avoided this bias by testing the algorithms on many different scenarios, altering the most crucial components of a stochastic and non-stationary multi armed bandit, while keeping the parameters that were tuned on a simple mid-case. For the mid-case we considered 5 arms, with initial arm probabilities of 0.8, 0.4, 0.3, 0.2, 0.1 for each arm respectively, with a circular shift of those at every 2000 time steps, in a total of 10000. After optimization, the parameters chosen for D-UCB were *ξ* = 0.22 and *γ* = 0.9999, for SW-UCB *ξ* = 0.3 and *τ* = 998, for Adapt-EvE *δ* = 0.13, *λ* = 40, *t_m_* = 50 and used UCBT as decision-makers and as meta-bandit. For KF-MANB *σ_ob_* = 0.2, *σ_tr_* = 0.01, initializing all means and variances to 0.5. For MLB *α_Q_* = 0.14, *α_m_* = 1/15, *α_ℓ_* = 1/350, *η* = 0.44. For MLB-KF we kept the parameters found for MLB and KF-MANB and only tuned *ϕ* = 1.5.

For problem set I, we considered that a global change point *cp,* may occur at every time step *t* with a constant probability *h* (i.e *p_t_*(*cp*) = *h*). When a change point occurs, all arm probabilities are then re-sampled from a uniform distribution in [0, 1]. For each subproblem (using only the set of probabilities generated for each arm with the above type of random walk), we run each algorithm for 20 sub-sessions, then regenerated another problem set with the same probability *h*, and repeated the procedure for 100 hyper-sessions. We then calculated the average total cumulative regret with the assumption that the results are representative due to central limit theorem. We increased *h* and repeated all the above, for *h* ∈ [2/10000,8/10000] with a step of 1/10000. Therefore, an average of 2 to 8 global change points occur (we have an horizon of 10000) for every step. As seen in Fig 1 (left), MLB exhibited the best performance for small *h*, KF-MANB demonstrated both robustness and low regret in all cases and MLB-KF inherited the best of these performances by following the regret dynamics of the most optimal algorithm between the two. In problem set II we explored cases where the difference ∆_*g*_ between the expected reward of the optimal action and the second best action varied. The best optimal arm always had a probability of 0.8, the second best 0.8−∆_*g*_, the third 0.8−2∆_*g*_ and so forth. For our experiments we begun with *#cp* = 1 (where *#cp* denotes the number of change points). This change point occurred at *t* = *T*/2, when the probability of the best arm dropped from 0.8 to 0.8-∆_*g*_, the probability of the second best dropped from 0.8-∆_*g*_ to 0.8-2∆_*g*_ (and so forth for other intermediate arms), while the probability of the worst arm increased from 0.8−4∆_*g*_ to 0.8. We altered ∆_*g*_ from 0.02 to 0.2 with a step size of 0.02 for each independent test case, while we simulated all algorithms on each case for 200 sessions, observing the average final cumulative regret. We then increased *#cp* by one, fixed the change point time steps to be evenly distributed over the timesteps, and repeated the procedure. In Fig.1 (middle) a visualization of the performance for each algorithm is presented, for different gaps ∆_*g*_ (rows), and *#cp* (columns). The results once again provided evidence that MLB-KF combines the behavior of both MLB and KF-MANB in a satisfactory manner, denoting the best performance on average. Finally, cases where the best optimal action becomes the less optimal, cases where the less optimal action becomes the optimal as also sequential alteration of those were tested in problem set III. MLB adapted greatly in Best-to-Worst scenarios and D-UCB in Worst-to-Best. However the overall average performance of MLB-KF was the empirically optimal as described in [11] (not shown here).

In order to highlight how parameters *v_t_*(*a*) change in time, a simple two-armed bandit case using KF-MANB, where *p_t_*(*a*_1_) was changing from 0.75 to 0.25 every 50 timesteps and *p_t_*(*a*_2_) = 1 − *p_t_*(*a*_1_), is also shown in Fig. 1 (right) for an average of 5 sessions. For the interval between timesteps 50-100, meta-parameter *v_t_*(*a*_1_) increases since the most chosen arm was *a*_2_. Even though the estimated reward rate of arm *a*_1_ in this interval differs from the true rate, a more precise estimation is not needed in order to make the most optimal decision. However, evidence about the arm’s reward rate in this interval is not provided, therefore *v_t_*(*a*_1_) increases, and then decreases as the arm is once again frequently chosen.

## 5 Conclusion and Discussion

In this work we presented empirical evidence that a hybrid algorithm which makes use of a classical computational neuroscience approach to properly tune exploration with meta-learning, combined with sibling Kalman filters proposed in Machine Learning to estimate each arm’s action value and add a measure of uncertainty as an exploration bonus, can lead to an adaptive learning strategy which efficiently manages the exploration-exploitation trade-off dilemma. However we still need to do the comparison more systematically for many different tasks (different numbers of arms, different stochasticity levels, different amplitudes between best and second-best arm, abrupt task shifts versus progressive drifts, different variation budget [13]). Nevertheless, such a bridge between the fields of computational neuroscience and machine learning seems promising.

Besides, each development of a new algorithm displaying efficient adaptation performance is an opportunity to sketch some new experimental predictions for neuroscience. Here, an interesting aspect of the hybrid algorithm is that it combines dynamic tuning of a global exploration rate *β_t_* through meta-learning and adaptive tuning of action specific learning rates *v_t_*(*a*) based on uncertainty variations in a Kalman filter. The resulting two timeseries may be less colinear than if they had been generated by the same method, which can be useful to generate regressors for model-based analyses of neurocience data. In addition, the resulting learning rates *v_t_*(*a*) may follow a different dynamics - briefly illustrated in Fig. 1 (right) - than other computational methods of learning rate studied in neuroscience bandit tasks, such as adaptations as a function of reward volatility [12]. Again, this may constitute an interesting experimental prediction to explore and compare in the future with human behavioral and neural data.

Finally, it is interesting to note recent progresses in machine learning using meta-learning combined with deep reinforcement learning [14]. Their deep learning approach is promising to not only dynamically tune some parameters but learn at a more global level the dynamics/statistics of the task. In addition, the method produces generalization to different but statistically related bandit tasks. All these results also suggest that a common ground between meta-learning and reinforcement learning can help find general methods to not only solve single-step bandit tasks, but also generalize to multi-step reinforcement leaning problems.

## Acknowledgements

This research work has been partially supported by the EU-funded Project BabyRobot (H2020-ICT-24-2015, grant agreement no. 687831), by the Agence Nationale de la Recherche (ANR-12-CORD-0030 Roboergosum Project and ANR-11-IDEX-0004-02 Sorbonne-Universités SU-15-R-PERSU-14 Robot Parallearning Project), and by Labex SMART (ANR-11-LABX-65 Online Budgeted Learning Project).

## References

[1] A. Garivier and E. Moulines, “On upper-confidence bound policies for non-stationary bandit problems,” Algorithmic Learning Theory, pp. 174–188, 2011.

[2] O.-C. Granmo and S. Berg, Solving Non-Stationary Bandit Problems by Random Sampling from Sibling Kalman Filters. Berlin, Heidelberg: Springer Berlin Heidelberg, 2010, pp. 199–208.

[3] N. D. Daw, J. P. O’Doherty, P. Dayan, B. Seymour, and R. J. Dolan, “Cortical substrates for exploratory decisions in humans,” Nature, vol. 441, no. 7095, pp. 876–879, 2006.

[4] R. C. Wilson, M. R. Nassar, and J. I. Gold, “Bayesian online learning of the hazard rate in change-point problems,” Neural computation, vol. 22, no. 9, pp. 2452–2476, 2010.

[5] P. Auer, N. Cesa-Bianchi, and P. Fischer, “Finite-time analysis of the multiarmed bandit problem,” Machine learning, vol. 47, no. 2-3, pp. 235–256, 2002.

[6] M. J. Frank, B. B. Doll, J. Oas-Terpstra, and F. Moreno, “Prefrontal and striatal dopaminergic genes predict individual differences in exploration and exploitation,” Nature neuroscience, vol. 12, no. 8, pp. 1062–1068, 2009.

[7] C. Hartland, S. Gelly, N. Baskiotis, O. Teytaud, and M. Sebag, “Multi-armed bandit, dynamic environments and meta-bandits,” in NIPS 2006 Workshop on Online trading between exploration and exploitation, Whistler, Canada, 2006.

[8] N. Schweighofer and K. Doya, “Meta-learning in reinforcement learning.” Neural Networks, vol. 16, no. 1, pp. 5–9, 2003.

[9] M. Khamassi, P Enel, P Dominey, and E. Procyk, “Medial prefrontal cortex and the adaptive regulation of reinforcement learning parameters,” Progress in Brain Research, vol. 202, pp. 441–464, 2013.

[10] M. Khamassi, G. Velentzas, T. Tsitsimis, and C. Tzafestas, “Active exploration and parameterized reinforcement learning applied to a simulated human-robot interaction task,” in IEEE Robotic Computing 2017, Taipei, Taiwan, 2017.

[11] G. Velentzas, C. Tzafestas, and M. Khamassi, “Bio-inspired meta-learning for active exploration during nonstationary multi-armed bandit tasks,” in IEEE Intelligent Systems Conference 2017, London, UK, 2017.

[12] T. E. Behrens, M. W. Woolrich, M. E. Walton, and M. F. Rushworth, “Learning the value of information in an uncertain world,” Nature neuroscience, vol. 10, no. 9, pp. 1214–1221, 2007.

[13] O. Besbes, Y. Gur, and A. Zeevi, “Optimal exploration-exploitation in a multi-armed-bandit problem with non-stationary rewards,” arXiv preprint arXiv:1405.3316, 2014.

[14] J. X. Wang, Z. Kurth-Nelson, D. Tirumala, H. Soyer, J. Z. Leibo, R. Munos, C. Blundell, D. Kumaran, and M. Botvinick, “Learning to reinforcement learn,” arXiv preprint arXiv:1611.05763, 2016.

